# EEG Functional Connectivity Reveals Accelerated Brain Aging in Young Adults with Cognitive Deficits and Mental Health Conditions

**DOI:** 10.64898/2026.09.17.752502

**Authors:** Chun-Ren Phang, Guo-Dong Miao, Lan-Tu Cai, Heng-Yi Rao, Ji-Xiang Du

## Abstract

The discrepancy between chronological age and predicted brain age, derived from neuroimaging data, serves as a biomarker for neurological health. However, the exact electroencephalography (EEG) mechanisms contributing to the variation of brain age are yet to be understood. This paper comprehensively investigated the relationship between brain age estimated from EEG-based functional connectivity and cognitive or mental manifestations of healthy adults. A mean absolute error (MAE) of 5.97 and R-squared (*R*^2^) of 0.85 were reported for training data, while an MAE of 10.73 and *R*^2^ of 0.56 were reported for the prediction data. Increased brain age was found to be negatively related to the scores of memory and attention tasks. Most young adults (19 of 26) suffering from mental symptoms such as addictions, depression, phobia, and anorexia nervosa demonstrated higher brain age than their chronological age. Alcohol consumption was also significantly correlated to the difference between brain and chronological ages. The functional connectivity of frontal and central motor regions was found to be most distinguishing between individuals with high and low brain ages. These brain regions were significantly correlated with brain age but not chronological age, suggesting a relatively stronger relationship with biological age. Ablation studies further confirmed the importance of the frontal brain and all bands EEG in the prediction of brain age. This work advances the understanding of EEG mechanisms contributing to brain age and could provide an interpretation for physiological or psychological conditions associated with brain age.

## 1 Introduction

The degeneration of the normal aging brain has been studied for years [1, 2]. Cognitive decline is one of the major symptoms of aging, accompanied by the loss of brain volume, especially in the frontal and parietal areas [3, 4]. Chronological age refers to the time metric calculated from the date of birth [5], while biological age is estimated based on health and physiological conditions [6]. Studies have indicated that chronological age might not always reflect the biological aging process [7, 8]. Cognitive decline was found to be more affected by biological age than chronological age [8].

Brain age, a biological age derived specifically from the brain, was introduced [9, 10]. Brain age is the machine-predicted age based on changes in brain structures or activity. The majority of recent research estimated brain age from magnetic resonance imaging (MRI) and functional MRI (fMRI) [11, 12]. Increased brain age was associated with diseases such as cognitive impairment, dementia, schizophrenia, Alzheimer’s disease, and diabetes mellitus [13, 14, 15, 11]. A mean absolute error (MAE) of 3.70 years was achieved with DeepBrainNet trained to predict brain age using 11,729 MRI images [14]. Grey matter volumetric maps were able to predict brain age with an MAE of 4.16 years [9]. An fMRI study predicted brain maturity using resting-state functional connectivity features from 238 scans [12]. Another study combined MRI structural and fMRI functional connectivity features for brain age prediction and achieved an MAE of 4.29 years [13]. A significantly increased brain age was found in individuals who are cognitively impaired.

Although MRI has achieved promising performance in predicting brain age, the prediction of brain age using electroencephalography (EEG) signals is rarely explored. To date, current EEG-based brain age prediction studies have focused mainly on sleep-related activities. The spectrogram and hypnogram computed from polysomnogram (PSG)-recorded EEG signals demonstrated a positive relationship between dementia severity and brain age predicted during sleep [15]. An MAE of 7.60 years was achieved by brain age predicted from EEG-derived time and frequency features during each of the five sleep stages [16]. This increased brain age was found to be associated with reduced life expectancy [17]. Amplitude, spectral, symmetry, and fractal features computed from 8-minute 32-channel EEG signals during relaxed, eyes-open conditions were able to predict brain age with an MAE of 6.89 years and an R-squared (*R*^2^) of 0.37 [18]. With just 2 minutes of data, a temporal convolutional network (TCN) was able to predict brain age from 21-channel EEG with an MAE of 6.60 years and an *R*^2^ of 0.73 [19].

Although functional connectivity extracted from EEG had been used to predict brain age [20], to our knowledge, the relationship between brain age and cognitive or mental manifestations in healthy adults has yet to be studied. Hence, the main objective of this study was to investigate how brain age derived from EEG functional connectivity relates to cognitive and mental manifestations in healthy adults. The major contributions of this study are demonstrating possible physiological or psychological conditions that are significantly associated with brain age. We also comprehensively studied how functional connectivity in different brain areas and EEG frequency bands were optimal in predicting brain age. The findings of this study are important for better understanding of the mechanisms underlying the variation between chronological and biological brain ages.

## 2 Methods

### 2.1 EEG Dataset

This study utilized the Leipzig Study for Mind-Body-Emotion Interactions (LEMON) open-source EEG dataset [21]. The LEMON dataset contains EEG, MRI, and fMRI recordings from young and elderly healthy adults with ages ranging from 20 to 77. To preserve anonymity, the publicly available dataset only provides ages in ranges of 5 years. This dataset is accompanied by rich physiological indicators including blood pressure, heart rate, smoking and drinking habits, disease history, and neurocognitive and behavioral functions, which are crucial to identifying the causes underlying the variation in brain age. The drinking habit was evaluated with the Alcohol Use Disorders Identification Test (AUDIT) [22], while the neurocognitive functions were mainly evaluated with the Wortschatztest verbal and crystallized intelligence test (WST) [23], Leistungsprüfsystem (LPS) fluid intelligence test [24], and Test of Attentional Performance (TAP) [25].

The 62-channel EEG recording was selected for this study. Out of the 216 EEG recordings, 5 were excluded due to incomplete data. The EEG signals were recorded with a BrainAmp MR plus amplifier and 62-channel (*N*_*ch*_ = 62) ActiCAP electrodes placed in accordance with the international 10–10 system. The reference electrode was placed at FCz, and the sampling rate was 2500 Hz. The impedance was maintained below 5 kΩ throughout the recording session, and a built-in 0.015 Hz to 1 kHz bandpass filter was applied. A total of 8 eye-closed (EC) and 8 eyes-open (EO) trials were recorded in a stratified manner, with each trial spanning 60 seconds.

### 2.2 Signal Preprocessing

To improve computational efficiency, the acquired time-series EEG signals were downsampled from 2500 Hz to 500 Hz, and the trials corresponding to EO and EC were extracted. No notch filter was used to ameliorate power line noise as the BrainAmp MR plus amplifier has built in function to eliminates noise originating from power supplies (visual inspection confirmed this) [26]. We only extracted the 30-second segments from the 15–45 seconds of the full 60-second segments to ameliorate the possible effects of transition periods between trials. These steps generated an *E × N*_*ch*_ *× T* matrix for each subject, where *E* = 16 represents the total number of EO and EC trials, *N*_*ch*_ = 62 is the number of EEG channels, and *T* = 15000 is the number of samples in 30 seconds. Subsequently, to study the effect of different frequency ranges, the EEG data were independently subjected to 7 bandpass filters (second-order, zero-phase Butterworth IIR filter), within the ranges of the wide band (0.1–100 Hz), delta band (0.1–3 Hz), theta band (4–7 Hz), alpha band (8–12 Hz), beta band (13–30 Hz), low gamma band (31–50 Hz), and high gamma band (51–100 Hz). The preprocessed data have the shape of *F × E × N*_*ch*_ *× T*, where *F* = 7 indicates the number of bandpassed frequencies.

### 2.3 Connectivity Features Extraction

The brain-wide functional connectivities were extracted from Pearson’s correlation. The Pearson’s correlation coefficient, *r*_*xy*_ *∈* [*−*1, 1], represents the weighted connectivity between pairwise EEG signals recorded from *x* and *y* channels. The connectivity strength was computed by dividing the covariance of signals *x* and *y* by the product of their standard deviations, as in Equation 1:

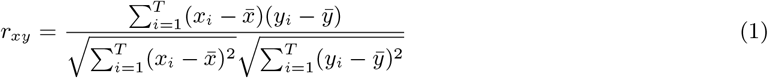

where 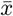 and 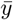 represent the means of EEG signals recorded from channels *x* and *y*, and *T* is the total number of time points. The aggregation of pairwise connections between all channels yielded a *N*_*ch*_ *× N*_*ch*_ connectivity matrix. The diagonal self-connections were set to zero. The connectivity matrices were independently extracted from the 7 frequency ranges, and trials from the same EO or EC tasks were averaged, generating a feature matrix of *F × N*_*ch*_ *× N*_*ch*_ *×* 2 for each individual.

To pinpoint the brain regions related to brain age, channel-wise degrees were computed from the connectivity matrices of specific frequency ranges. Degree is one of the main features of graph theory, which is important to reduce and characterize the high-dimensional complex brain connectivity network while retaining its meaningful information [27]. Degree, 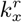, is the sum of connection strength for an individual channel *x*, and is used to measure the channel-wise local network integration, as shown in Equation 2:

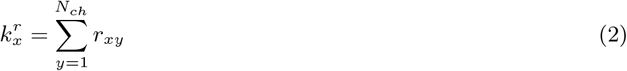

where *r*_*xy*_ is the weighted connectivity extracted using Pearson’s correlation between EEG signals recorded from channels *x* and *y*, and *N*_*ch*_ is the total number of channels. Similar to the connectivity matrix, degrees were extracted from each of the 7 frequency ranges, and the EO and EC trials were averaged independently. This produces a feature matrix of *F × N*_*ch*_ *×* 2 for each subject.

### 2.4 Regression Model

A multilayer perceptron with one input layer, one hidden layer, and one output layer was utilized as the regression model to predict brain age from functional connectivity and degree matrices. The size of the input layer depends on the size of the flattened feature matrices, while the hidden layer consisted of 64 nodes. An output node with a linear activation function was used for brain age prediction. A dropout rate of 0.5 was implemented in each layer to prevent overfitting. The model was trained with the ADAM optimizer with a learning rate of 0.001 [28] and a batch size of 64 for 500 epochs. The loss function used to tune the model weights was the mean absolute error (MAE), as in Equation 3:

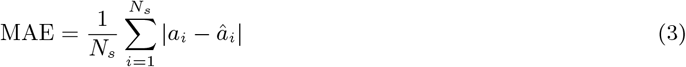

where *a*_*i*_ is the chronological age, *â*_*i*_ is the predicted brain age, and *N*_*s*_ = 211 is the number of subjects. As the chronological ages are in ranges of 5 years, the predicted brain ages were rounded to the nearest multiple of 5 before computing the loss.

Since Pearson’s correlation measures undirected pairwise connections (*x* to *y* connection and *y* to *x* connection are the same), the duplicate connections were removed. Both connectivity and degree feature matrices were flattened before feeding them into the regression model. We selected the all bands features and combined the EO and EC tasks, yielding input shapes of 3782 *×* 7 and 124 *×* 7 for connectivity and degree features, respectively.

### 2.5 Model Validation

The performance of the regression model was evaluated using a ten-fold cross-validation method, where in each fold, 90% of the data were used to train the model, and 10% were used for performance evaluation. To prevent data leakage and overfitting, all the 211 data points were treated as independent samples without further partitioning, where the testing data did not contain features from subjects in the training data. The model performance was evaluated using two metrics, namely MAE and R-squared (*R*^2^). Equation 4 demonstrates the computation of *R*^2^:

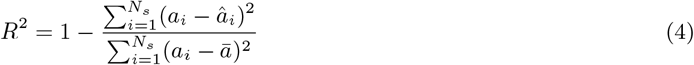

where *a*_*i*_ is the chronological age, *â*_*i*_ is the predicted brain age, and *N*_*s*_ is the number of subjects. A preliminary analysis found that degree features poorly predict brain age, hence, the subsequent ablation studies omitted the degree features.

### 2.6 Ablation Studies

The ablation studies were based only on functional connectivity features, as the degree features had low predictability. Four ablation protocols were implemented on the feature set, including the ablation of frequency, brain regions, recording channels, and tasks. Instead of using the connectivity features extracted from the all frequency bands (7 × 64 × 64 × 2), we investigated the performance of features extracted from specific EEG frequency bands (1 × 64 × 64 × 2). The brain regions were grouped based on lobes, such as anterior (*N*_*ch*_ = 5), frontal (*N*_*ch*_ = 19), central (*N*_*ch*_ = 14), parietal (*N*_*ch*_ = 16), temporal (*N*_*ch*_ = 4), and occipital (*N*_*ch*_ = 3) regions. The MAE and *R*^2^ of the combination of anterior and frontal (*N*_*ch*_ = 24) regions were also evaluated. For the ablation of recording channels, the performance of specific combinations of channel labels was studied, including FP+F (*N*_*ch*_ = 5), FP+C (*N*_*ch*_ = 5), F+C (*N*_*ch*_ = 6), FP+F+C (*N*_*ch*_ = 8), F+C+P (*N*_*ch*_ = 9), and FP+F+C+P (*N*_*ch*_ = 11). Lastly, we also investigated the effect of eye conditions on the regression performance by comparing the evaluation metrics computed from EO or EC data.

### 2.7 Statistical Analysis

Due to the bounded nature of the data and the ordinal nature of variables, group comparisons between young and older adults were performed using the nonparametric Mann-Whitney U test [29]. This test does not assume a specific distribution and is suitable for the types of data analyzed. All correlation values reported in this study represent Pearson’s correlation coefficients (PCC). For both statistical comparisons and correlation analyses, p-values were Bonferroni-corrected to account for multiple comparisons across feature numbers [30]. The mean, standard deviation (SD), and interquartile range (IQR) were reported.

## 3 Results

### 3.1 Participant Demographics

Out of 211 individuals, 141 (44 female, 97 male) were young adults aged 35 or below, while 70 (35 female, 35 male) were older adults aged 55 or above. The mean age of young adults was 22.87, with a standard deviation (SD) of 3.33 and an interquartile range (IQR) of 5.00; the mean age of older individuals was 65.14, with an SD of 5.00 and an IQR of 10.00. Significant differences in blood pressure were found between the two age groups, where young individuals had a lower mean systolic pressure of 122.36 (SD: 11.86, IQR: 18.00) and a mean diastolic blood pressure of 74.74 (SD: 8.10, IQR: 11.00) compared to the systolic pressure of 154.84 (SD: 19.85, IQR: 23.00) and diastolic pressure of 89.50 (SD: 10.86, IQR: 14.50) in older individuals. Young adults (5.65, SD: 4.04, IQR: 4.00) also consumed more alcohol than older individuals (3.09, SD: 2.53, IQR: 2.00), based on the difference in AUDIT alcohol use scores. In terms of cognition test scores, young adults demonstrated significantly higher WST, LPS, and TAP-memory correct rates and faster reaction times in TAP tasks compared to older individuals. These results are summarized in Table 1.

**Table 1:** Demographic characteristics, physiological conditions, and cognitive test scores [MEAN (SD, IQR)] in the subject data from the MPI-Leipzig Mind-Brain-Body study (*N* = 211). Mind-Brain-Body study (*N* = 211). BPM: Beats per Minute; AUDIT: Alcohol Use Disorders Identification Test [22]; WST: Wortschatztest verbal and crystallized intelligence test [23]; LPS: Leistungsprufsystem fluid intelligence test [24]; TAP: Test of Attentional Performance [25].

|  | Young (age<=35) | Old (age>=55) | p-value |
| --- | --- | --- | --- |
| Age (years) | 22.87 (3.33, 5.00) | 65.14 (5.00, 10.00) | **$p < 0.01$ |
| Gender | 44 F, 97 M | 35 F, 35 M | $p > 0.05$ |
| Systole Blood Pressure (mmHg) | 122.36 (11.86, 18.00) | 154.84 (19.85, 23.00) | **$p < 0.01$ |
| Diastole Blood Pressure (mmHg) | 74.74 (8.10, 11.00) | 89.50 (10.86, 14.50) | **$p < 0.01$ |
| Heart Rate (BPM) | 65.52 (9.87, 13.00) | 67.24 (9.39, 11.75) | $p > 0.05$ |
| Smoking (Non-smoker=1, Occasional Smoker=2, Smoker=3) | 1.28 (0.61, 0.00) | 1.14 (0.46, 0.00) | $p > 0.05$ |
| Hamilton Scale | 2.57 (2.62, 4.00) | 2.10 (2.49, 3.00) | $p > 0.05$ |
| AUDIT | 5.65 (4.04, 4.00) | 3.09 (2.53, 2.00) | **$p < 0.01$ |
| Standard Alcoholunits Last 28 days | 18.30 (18.14, 22.00) | 17.542 (25.43, 19.13) | $p > 0.05$ |
| WST Correct | 33.00 (2.94, 3.00) | 33.29 (2.75, 3.00) | $p > 0.05$ |
| LPS Correct | 21.21 (3.46, 4.00) | 15.61 (2.91, 4.75) | **$p < 0.01$ |
| TAP Compatible Reaction Time (s) | 409.30 (77.58, 75.00) | 522.81 (115.38, 123.25) | **$p < 0.01$ |
| TAP Compatible Incorrect | 1.07 (2.73, 1.00) | 1.64 (3.88, 2.00) | $p > 0.05$ |
| TAP Incompatible Reaction Time (s) | 436.96 (68.07, 87.00) | 587.17 (125.12, 125.75) | **$p < 0.01$ |
| TAP Incompatible Incorrect | 1.61 (3.25, 2.00) | 2.99 (4.75, 3.75) | $p > 0.05$ |
| TAP Memory Correct Reaction Time (s) | 551.37 (144.40, 168.00) | 627.90 (186.15, 223.75) | *$p < 0.05$ |
| TAP Memory Correct | 85.88 (13.78, 15.97) | 65.92 (23.84, 36.96) | **$p < 0.01$ |
| TAP Memory Incorrect | 6.89 (8.25, 11.76) | 17.55 (15.04, 22.32) | **$p < 0.01$ |
| TAP Memory Missed | 7.24 (9.32, 10.00) | 16.53 (14.68, 18.85) | **$p < 0.01$ |

### 3.2 Relationship between Chronological and Brain Age

Figure 1 visualizes the relationship between chronological age and machine-predicted brain age. A preliminary analysis found that degree features poorly predict brain age, hence, this section elaborate the prediction of brain age using functional connectivity features. The Pearson’s correlation coefficients (PCC) between both ages were 0.92 for training data and 0.75 for testing data. The mean absolute errors (MAE) were 5.97 and 10.73 for training and testing data, respectively, while the R-squared (R^2^) values were 0.85 and 0.56 for training and testing data, respectively. Out of the 141 young adults, 1 had matched brain age, 42 had low brain age, and 98 had high brain age. For the old adults, 4 had matched brain age, 56 had low brain age, and 10 had high brain age. To enhance distinctiveness, we specifically classified very high brain age (VHBA) as [brain age *>* chronological age +10] and very low brain age (VLBA) as [brain age *<* chronological age −10] in the subsequent analysis. This yielded 8 young adults with VLBA, 41 young adults with VHBA, 40 old adults with VLBA, and 1 old adult with VHBA. Since the numbers of young adults with VLBA and old adults with VHBA were limited, these groups were not included in the subsequent analysis to ensure statistical robustness.

**Figure 1:**
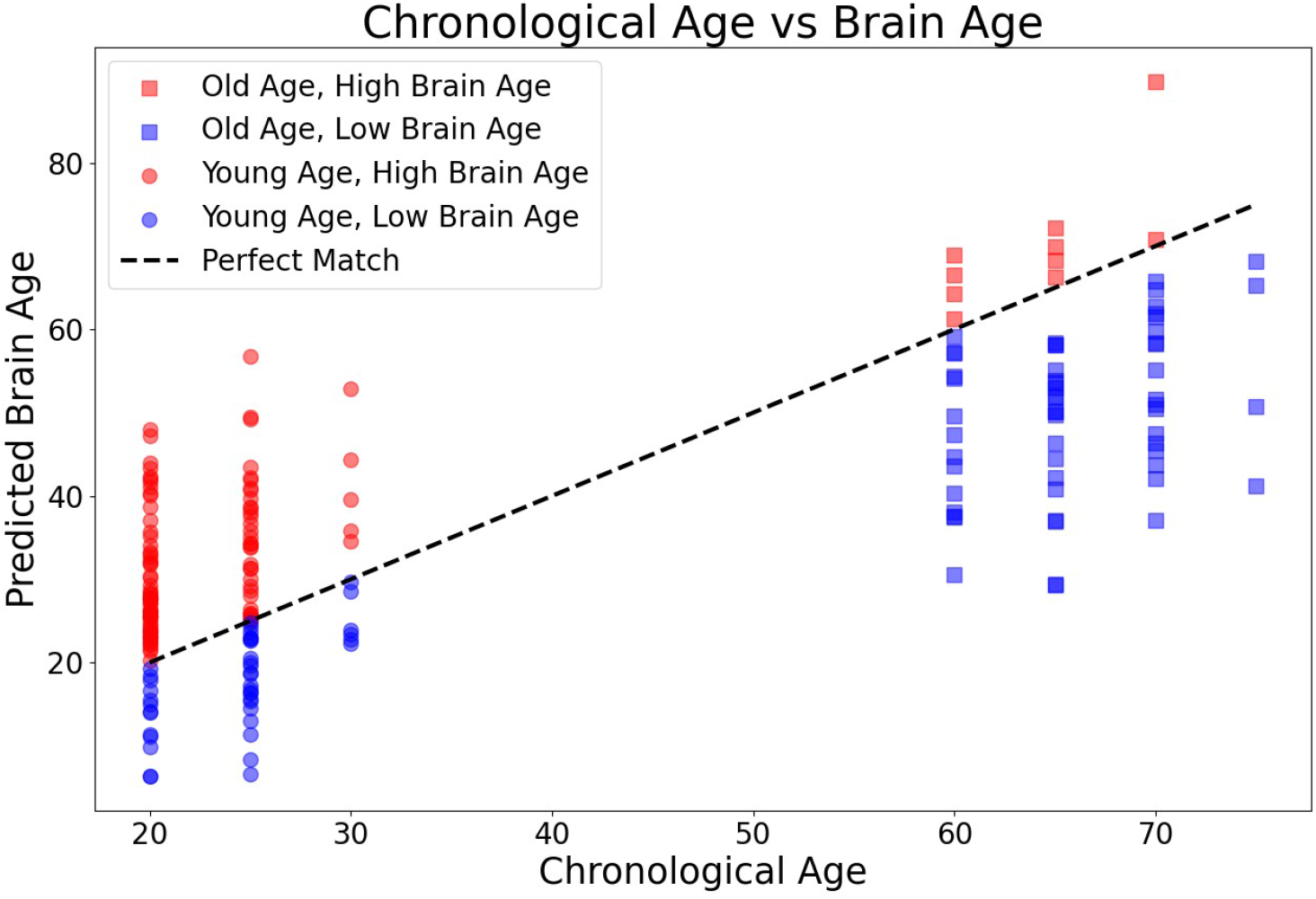
The linear fit between chronological age and machine predicted brain age. Ten old and 98 young individuals have high predicted age (high brain age), while 56 old and 42 young individuals have low predicted age (low brain age).

#### 3.2.1 Brain Age and Cognitive Scores

Significant differences in cognitive scores were found between individuals with VLBA or VHBA, as shown in Table 2. Young individuals with VHBA reported a correct rate of 81.31% (SD: 15.81, IQR: 22.75) in TAP memory tasks, which was significantly lower (\**p <* 0.05) than the correct rate of 87.75% (SD: 12.37, IQR: 12.96) in young baseline individuals without VHBA. In the TAP incompatible task, young adults with VHBA reported significantly higher (\*\**p <* 0.01) incorrect rates of 2.12% (SD: 3.24, IQR: 1.00) compared to the incorrect rate of 1.40% (SD: 3.24, IQR: 2.00) in baseline young individuals. Counterintuitively, baseline young individuals demonstrated significantly higher (\**p <* 0.05) incorrect rates of 1.32% (SD: 3.17, IQR: 1.25) in the TAP compatible task compared to the incorrect rate of 0.46% (SD: 0.80, IQR: 1.00) in VHBA young adults. On the other hand, the WST correct rate of old age baseline individuals was marginally significantly lower (p=0.05) than that of old individuals with VLBA, with reported correct rates of 32.50% (SD: 2.64, IQR: 3.00) and 33.88% (SD: 2.69, IQR: 3.00), respectively.

**Table 2:** Cognitive test scores [MEAN (SD, IQR)] of individuals with very high brain age compared to young age and individuals with very low brain age compared to old age.

|  | Very High Brain Age | Young Age | p-value |
| --- | --- | --- | --- |
| TAP Memory Correct (%) | 81.31 (15.81, 22.75) | 87.75 (12.37, 12.96) | $*p < 0.05$ |
| TAP Incompatible Incorrect (%) | 2.12 (3.24, 1.00) | 1.40 (3.24, 2.00) | $**p < 0.01$ |
| TAP Compatible Incorrect (%) | 0.46 (0.80, 1.00) | 1.32 (3.17, 1.25) | $*p < 0.05$ |
|  | Very Low Brain Age | Old Age | p-value |
| WST Correct (%) | 33.88 (2.69, 3.00) | 32.50 (2.64, 3.00) | $p = 0.05$ |

#### 3.2.2 Brain Age and Mental-Related Conditions

From the original dataset of 211 subjects, 40 subjects were diagnosed with mental disorders based on structured clinical interviews according to the Diagnostic and Statistical Manual of Mental Disorders version 4 (DSM-IV) [31]. These results are tabulated in Table 3. The diagnosed disorders can be classified into three groups, including alcohol or cannabis addictions, psychological symptoms, and physical abuse. Fourteen of the diagnosed subjects were chronologically old individuals (age*>*=55), where 2 of them who suffered from psychological symptoms including depression and dysthymia had high brain age. Of the 26 chronologically young individuals (age*<*=35), 19 of them who had high brain age were either isolated or combined addicted to alcohol (N=6) or cannabis (N=4), abused (N=1), diagnosed with depression (N=4), phobia (N=4), bipolar disorder (N=1), panic disorder (N=1), obsessive-compulsive disorder (OCD) (N=1), tic disorder (N=1), or anorexia nervosa (N=1). We also found 8 young VHBA adults who were isolated or combined suffering from addiction to alcohol (N=4) or cannabis (N=4), phobia (N=2), depression (N=1), or anorexia nervosa (N=1). These results suggest that alcohol or cannabis consumption could be related to brain aging in young adults. Further analysis also found a significant (\*\**p <* 0.01) positive correlation of 0.35 between Δage (brain age minus chronological age) and scores of Alcohol Use Disorders Identification Test (AUDIT) [22], visualized in Figure 2.

**Table 3:** Number of individuals with high brain age who reported mental-related conditions. Most of the individuals suffering from mental-related conditions are young adults with high brain age.

|  | Number of individual | Addiction | Psychological | Abused |
| --- | --- | --- | --- | --- |
| High Brain Age in Old Age | 2 in 14 | 0 | 2 | 0 |
| High Brain Age in Young Age | 19 in 26 | 7 | 11 | 1 |
| Very High Brain Age ($>10$) in Young Age | 8 in 26 | 4 | 4 | 0 |

**Figure 2:**
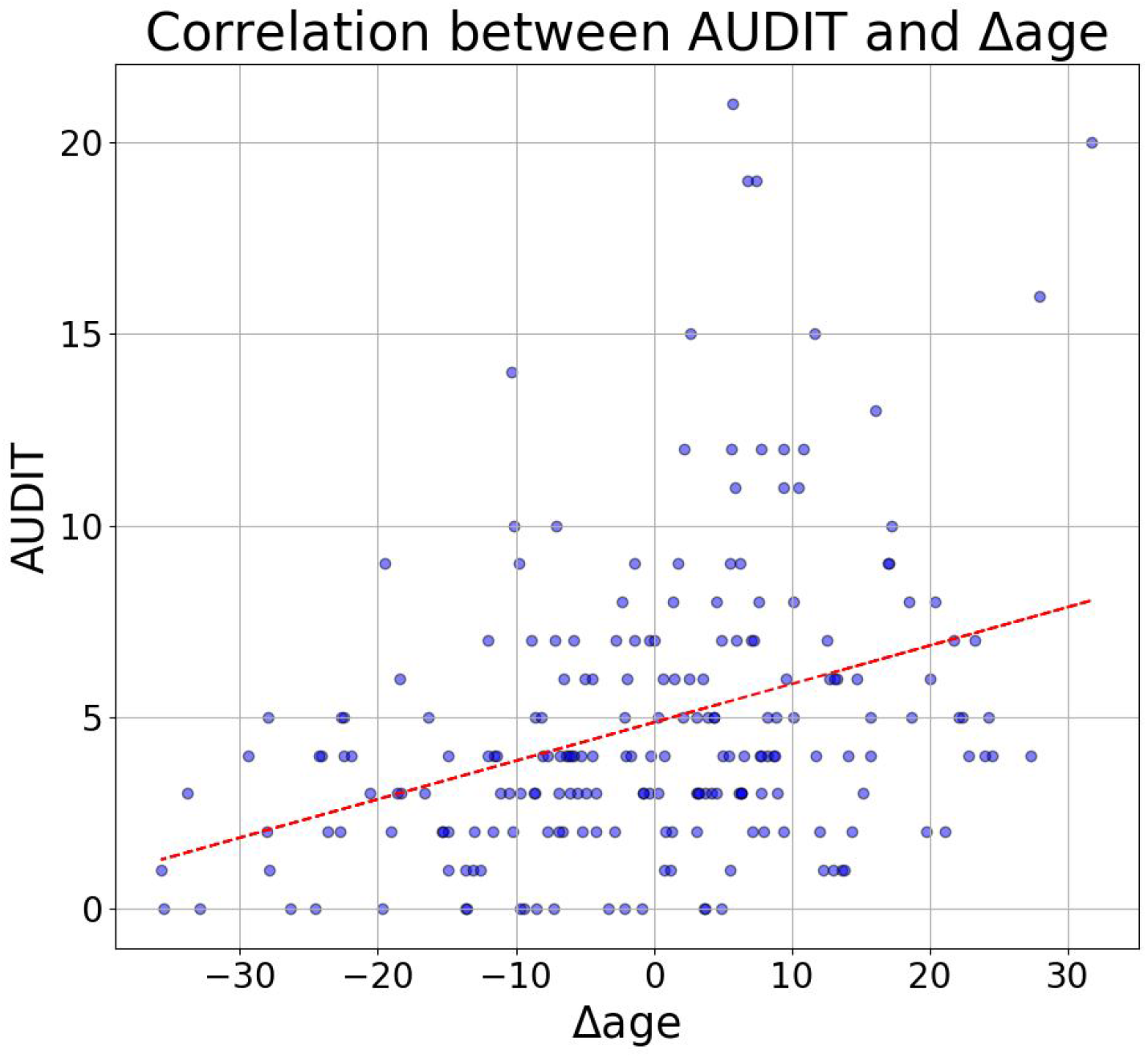
The correlation (*PCC* = 0.35, \*\**p <* 0.01) between Alcohol Use Disorders Identification Test (AUDIT) and Δage (brain age minus chronological age).

### 3.3 Functional Connectivities Related to Very High and Very Low Brain Age

We further analyzed the channel-based pairwise functional connectivity strength in VLBA and VHBA subjects. Table 4 summarizes the significant connection differences between old individuals with and without VLBA. The significant differences were only observed during the 1-minute opened eyes condition but not during closed eyes. VLBA subjects demonstrated 11 lower intralobular frontal connectivities between frontopolar (FP), anterior frontal (AF), and frontal (F) brain regions in delta, theta, alpha, and wide-range EEG frequencies. Seven were intrahemispheric connections (wide FP1-AF7, delta FP1-AF7, delta AF4-AF8, theta FP1-AF7, theta FP2-AF8, theta AF4-AF8, alpha FP1-AF7), while four were interhemispheric connections including midline connections (delta FP2-AFz, alpha AFz-AF7, theta FP2-AFz, theta FP2-F3).

**Table 4:** The significant different of functional connectivities [MEAN (SD, IQR)] between old individual with and without very low brain age during opened eyes condition (*p <* 0.05; effect size*>* 0.3). The majority of the significant brain activities are delta, theta, and alpha frequencies in prefrontal regions.

|  | Very Low Brain Age | Old Age | Difference |
| --- | --- | --- | --- |
| Wide FP1-AF7 | 0.84 (0.11, 0.16) | 0.91 (0.07, 0.06) | -0.07 |
| Delta FP1-AF7 | 0.88 (0.15, 0.09) | 0.94 (0.05, 0.04) | -0.06 |
| Delta FP2-AFz | 0.81 (0.19, 0.18) | 0.93 (0.05, 0.06) | -0.12 |
| Delta AF4-AF8 | 0.79 (0.15, 0.21) | 0.90 (0.05, 0.07) | -0.11 |
| Theta FP1-AF7 | 0.96 (0.03, 0.04) | 0.98 (0.03, 0.03) | -0.02 |
| Theta FP2-F3 | 0.65 (0.16, 0.19) | 0.75 (0.13, 0.17) | -0.10 |
| Theta FP2-AFz | 0.88 (0.15, 0.09) | 0.94 (0.05, 0.04) | -0.06 |
| Theta FP2-AF8 | 0.89 (0.14, 0.07) | 0.95 (0.03, 0.05) | -0.06 |
| Theta AF4-AF8 | 0.90 (0.06, 0.10) | 0.94 (0.05, 0.04) | -0.04 |
| Alpha FP1-AF7 | 0.91 (0.04, 0.05) | 0.94 (0.04, 0.03) | -0.03 |
| Alpha AFz-AF7 | 0.73 (0.07, 0.09) | 0.79 (0.09, 0.09) | -0.06 |

Table 5 compares the significant connection differences between young individuals with and without VHBA. In line with VLBA groups, significant differences were only observed during the opened eyes condition. Sixteen significant connections in central (C), frontal (F), frontocentral (FC), and parietocentral (CP) brain areas were found to be lower in VHBA young subjects. Partially similar to VLBA old subjects, the differences were present in theta, beta, low gamma, and wide-range EEG frequencies. Four were intrahemispheric connections (wide FC2-C4, wide FC2-C6, theta FC2-C8, low gamma FC1-C3), while twelve were interhemispheric connections (wide FC2-C3, wide FC2-C7, theta FC2-FC1, theta FC2-C3, theta FC2-C5, theta FC2-C1, beta FC2-C1, low gamma FC2-CP1, low gamma FC2-C1, low gamma FC1-C4, low gamma C2-F3, low gamma C2-F1).

**Table 5:** The significant different of functional connectivities [MEAN (SD, IQR)] between young individual with and without very high brain age during opened eyes condition (*p <* 0.05; effect size*>* 0.5). The majority of the significant brain activities are theta and low gamma frequencies in frontocentral regions.

|  | Very High Brain Age | Young Age | Difference |
| --- | --- | --- | --- |
| Wide FC2-C4 | 0.05 (0.17, 0.19) | 0.19 (0.15, 0.17) | -0.14 |
| Wide FC2-C3 | -0.01 (0.18, 0.20) | 0.15 (0.16, 0.19) | -0.16 |
| Wide FC2-C6 | 0.10 (0.15, 0.14) | 0.22 (0.14, 0.16) | -0.12 |
| Wide FC2-C7 | 0.13 (0.14, 0.19) | 0.25 (0.14, 0.17) | -0.12 |
| Theta FC2-FC1 | -0.19 (0.16, 0.21) | 0.01 (0.23, 0.30) | -0.20 |
| Theta FC2-C8 | 0.21 (0.14, 0.21) | 0.33 (0.13, 0.16) | -0.12 |
| Theta FC2-C3 | 0.05 (0.16, 0.21) | 0.22 (0.18, 0.26) | -0.17 |
| Theta FC2-C5 | 0.11 (0.17, 0.21) | 0.27 (0.16, 0.21) | -0.16 |
| Theta FC2-C1 | 0.02 (0.17, 0.21) | 0.20 (0.20, 0.27) | -0.19 |
| Beta FC2-C1 | -0.16 (0.14, 0.21) | -0.02 (0.15, 0.19) | -0.14 |
| Low Gamma FC2-CP1 | 0.05 (0.16, 0.18) | 0.18 (0.13, 0.14) | -0.13 |
| Low Gamma FC2-C1 | -0.06 (0.16, 0.22) | 0.09 (0.14, 0.14) | -0.15 |
| Low Gamma FC1-C4 | -0.08 (0.17, 0.28) | 0.07 (0.13, 0.14) | -0.15 |
| Low Gamma FC1-C3 | -0.04 (0.14, 0.18) | 0.09 (0.11, 0.13) | -0.13 |
| Low Gamma C2-F3 | -0.16 (0.13, 0.19) | -0.04 (0.12, 0.13) | -0.12 |
| Low Gamma C2-F1 | -0.18 (0.16, 0.19) | -0.04 (0.12, 0.16) | -0.14 |

### 3.4 Degrees Features Related to Very High and Very Low Brain Age

Figure 3 displays scalp topographical maps highlighting brain regions associated with VLBA and VHBA. Older individuals with VLBA exhibited elevated degrees across delta, theta, alpha, beta, and wide-band frequencies compared to baseline young controls, as shown in Figure 3(a). These differences were predominantly localized to left-hemisphere regions, including frontal, central, temporal, and occipital areas. Notably, these effects were specific to the eyes-open condition, with no significant findings observed during eyes-closed recording. Young adults with VHBA showed reduced degrees in most frequency bands during both eyes-open and eyes-closed conditions, except for delta and beta band activity during eyes-closed (Figure 3b). The bilateral hemispheric reductions were particularly evident in wide-band temporal regions, frontal lobe delta, theta, alpha, and beta activity, and gamma-band frontal and temporal connectivity degree.

**Figure 3:**
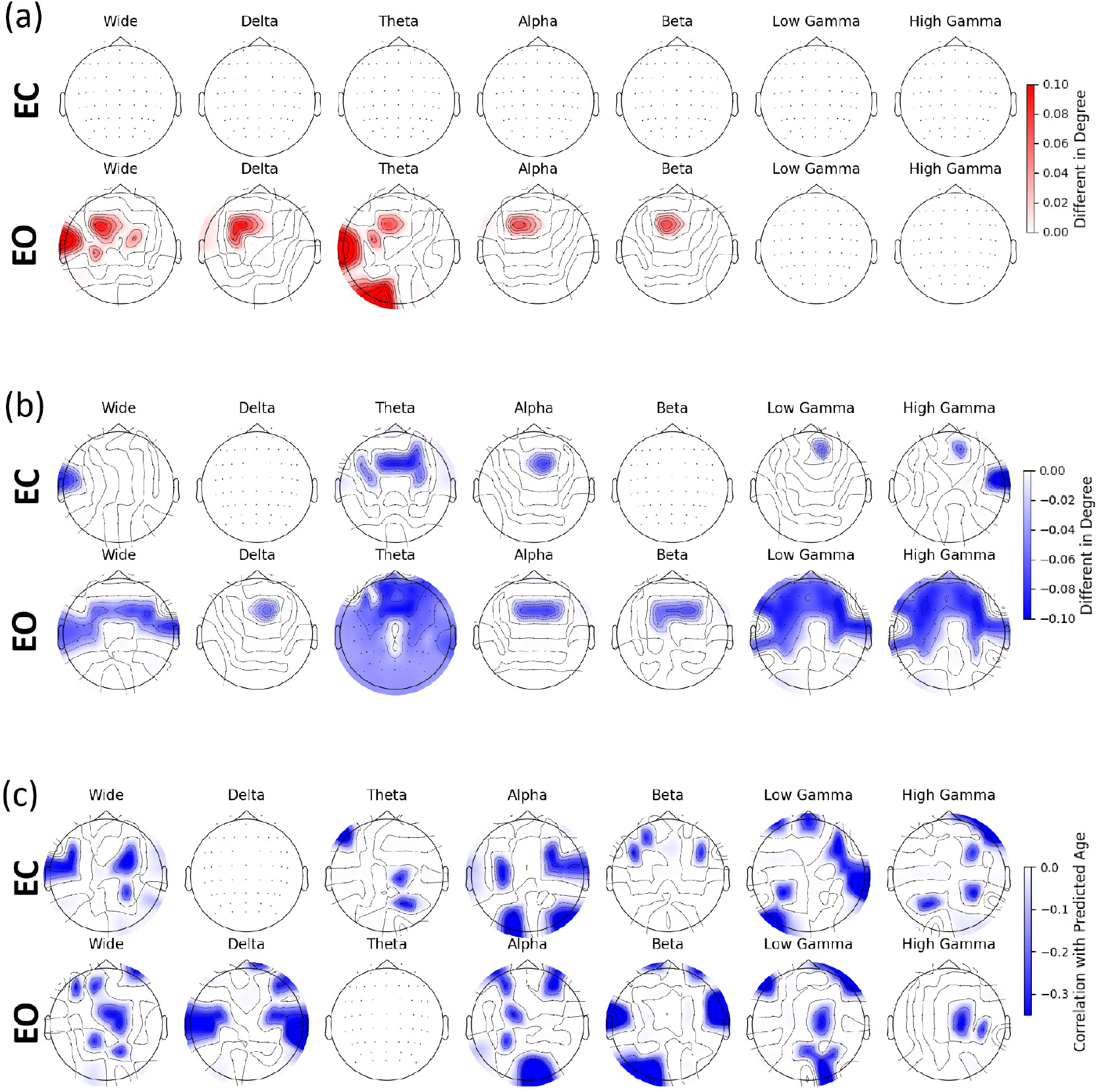
The scalp topomaps of (a) the different \**p <* 0.05 in degrees between old individuals with and without very low brain age; (b) the different \**p <* 0.05 in degrees between young individuals with and without very high brain age; (c) the significant correlation of degrees (\**p <* 0.05) to brain age which are not significantly correlated to chronological age.

To identify brain-age-specific regions, we examined degree features that correlated significantly with predicted brain age but not chronological age. Multiple regions across frontal, central, temporal, parietal, and occipital cortices met these criteria across seven frequency bands, as illustrated in Figure 3(c). By intersecting these correlation patterns with the group difference maps (Figures 3(a) & 3(b)), we identified distinct spatial profiles where VLBA was characterized by increased connectivity in bilateral frontocentral regions, while VHBA primarily showed enhancements in left temporal and frontal areas (Figure 4).

**Figure 4:**
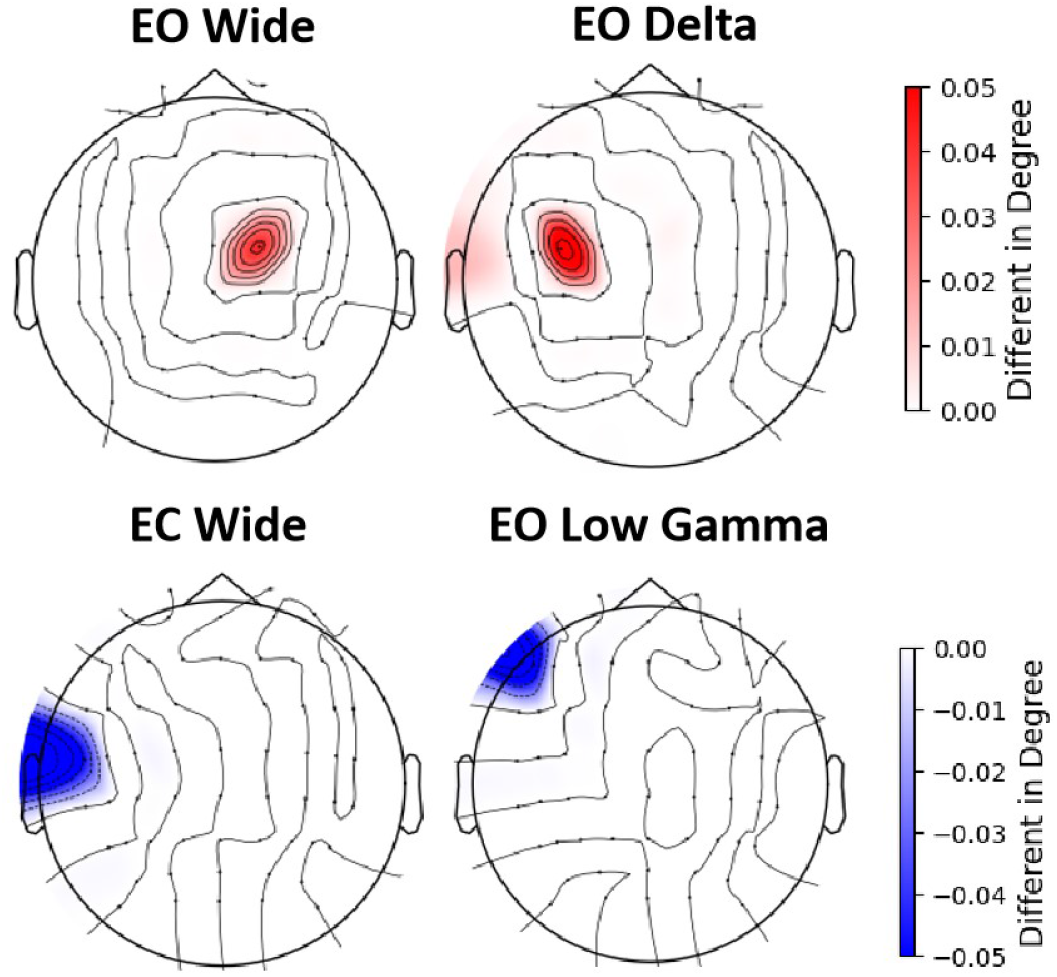
The scalp topomaps demonstrate the intersection between very low brain age (upper panel) and very high brain age (lower panel) regions and the brain regions significantly correlated to brain age.

### 3.5 Brain Age Predictions with Different EEG Activities

This section demonstrates the performance of brain age prediction using different EEG activities. Figure 5(a) investigates the effect of eight EEG frequency ranges on prediction performance. All bands represent the concatenation of connectivity matrices extracted from delta, theta, alpha, beta, low and high gamma bands, yielding an *N*_*ch*_ *× N*_*ch*_ *× N*_*task*_ *× N*_*band*_ feature dimension. The reported mean absolute error (MAE) and R-squared (R^2^) for wide-bands (0.1-100 Hz filtered EEG signals) were 14.33 and 0.21, respectively. The lowest MAE of 10.73 and highest R^2^ of 0.56 were found in the all bands connectivity features. Conversely, the highest MAE of 15.66 and lowest R^2^ of 0.09 were found in the delta band connectivity feature. MAEs of 14.16, 13.74, 13.18, 12.79, and 13.24, and R^2^ values of 0.27, 0.30, 0.35, 0.35, and 0.31 were reported for theta, alpha, beta, low gamma, and high gamma bands, respectively.

**Figure 5:**
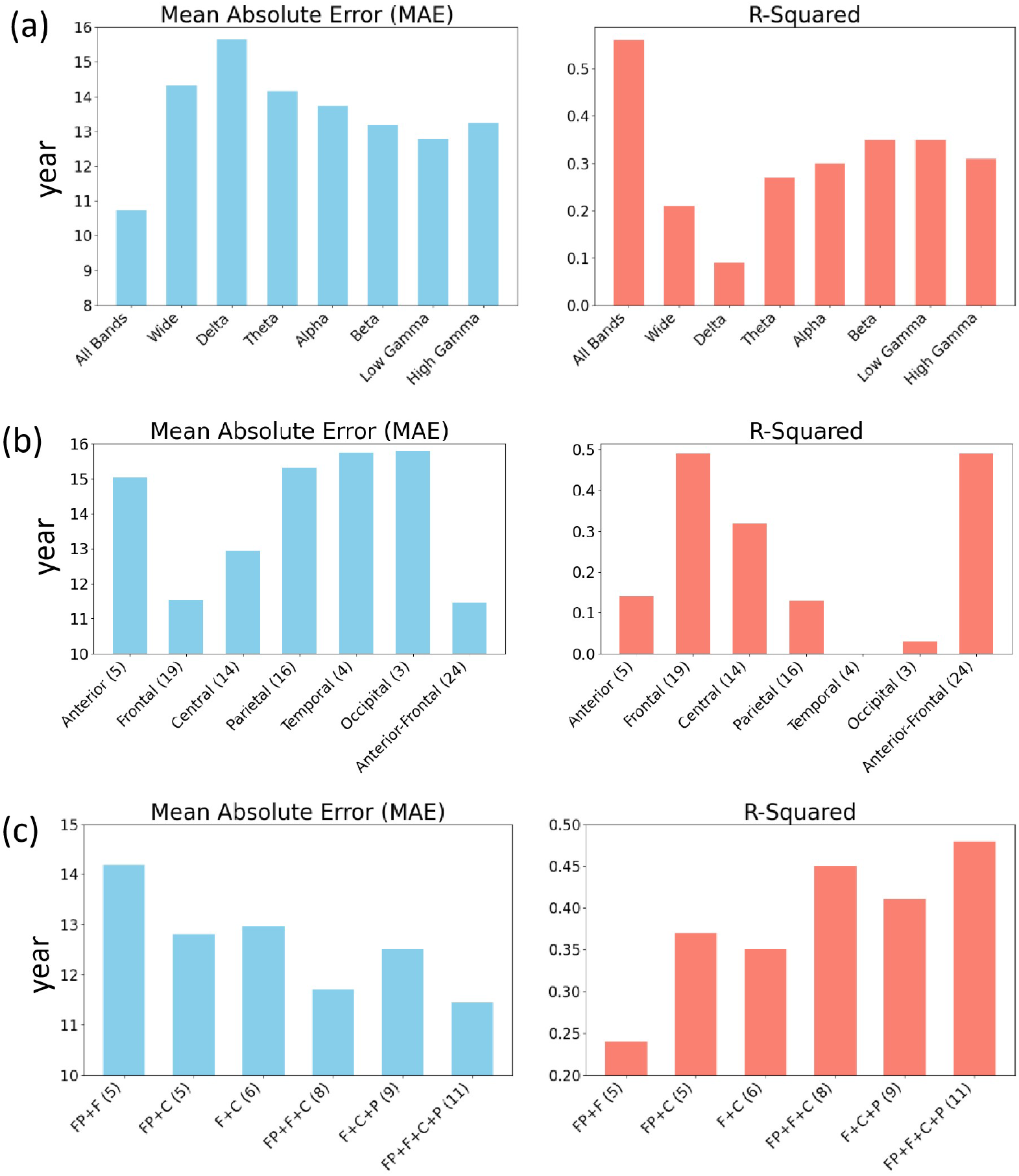
The bar graphs visualize the mean absolute error (MAE) and R-square (R^2^) of regression models trained and tested with different (a) frequency bands, (b) brain regions, and (c) channel locations. Brackets indicate the number of channel used.

Figure 5(b) examines the effect of brain regions on brain age prediction performance. The functional connectivity features were clustered into anterior (*N*_*ch*_ = 5), frontal (*N*_*ch*_ = 19), central (*N*_*ch*_ = 14), parietal (*N*_*ch*_ = 16), temporal (*N*_*ch*_ = 4), and occipital (*N*_*ch*_ = 3) regions. The performance of combined anterior and frontal regions (*N*_*ch*_ = 24) was also studied. The anterior-frontal regions (MAE=11.46; R^2^=0.49) achieved the best performance, followed by frontal (MAE=11.52; R^2^=0.49) and central (MAE=12.95; R^2^=0.32) regions. Suboptimal performances were observed in anterior (MAE=15.04; R^2^=0.14), parietal (MAE=15.31; R^2^=0.13), temporal (MAE=15.74; R^2^=0.00), and occipital (MAE=15.78; R^2^=0.03) lobes.

To further identify the importance of EEG recording channels, we analyzed the prediction performance of various channel combinations, including FP+F (*N*_*ch*_ = 5), FP+C (*N*_*ch*_ = 5), F+C (*N*_*ch*_ = 6), FP+F+C (*N*_*ch*_ = 8), F+C+P (*N*_*ch*_ = 9), and FP+F+C+P (*N*_*ch*_ = 11). The results indicate that using only frontal channels (FP+F) was insufficient for brain age prediction, as it yielded the highest MAE of 14.18 and lowest R^2^ of 0.24. The FP+C, F+C, and F+C+P combinations showed similar performance, with MAEs of 12.81, 12.96, and 12.50, and R^2^ values of 0.37, 0.35, and 0.41, respectively. Higher performance was demonstrated by FP+F+C and FP+F+C+P, where adding two parietal channels to FP+F+C (creating FP+F+C+P) decreased the MAE from 11.71 to 11.44 and increased the R^2^ from 0.45 to 0.48. These results are visualized in Figure 5(c).

Based on these findings, FP+F+C appears to be the optimal channel combination, achieving a balance between prediction performance and channel count. Figure 6 compares the MAE and R^2^ during eyes-open (EO) and eyes-closed (EC) conditions. The EC condition (MAE=11.99; R^2^=0.43) performed similarly to the combined EC and EO conditions (MAE=11.71; R^2^=0.45), while both outperformed the EO condition alone (MAE=13.59; R^2^=0.30). These results suggest that functional connectivity extracted from frontopolar, frontal, and central EEG channels during the eyes-closed condition provides optimal brain age prediction.

**Figure 6:**
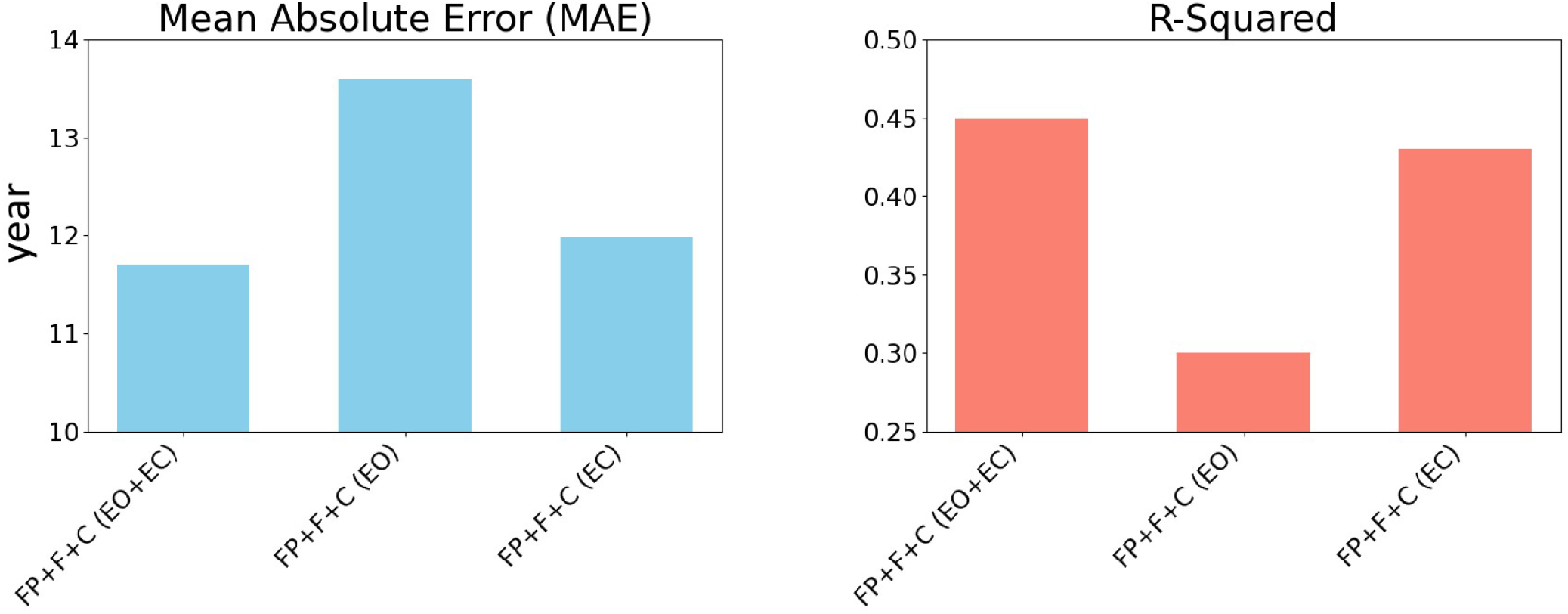
The bar graphs visualize the mean absolute error (MAE) and R-square (R^2^) of FP+F+C regression models trained and tested with opened and closed eyes conditions.

## 4 Discussion and Conclusion

The results indicate that VHBA young adults have significantly poorer working memory capacity in the TAP memory task. While tasked with remembering the second last number of a sequence, VHBA young adults demonstrate lower correct response rates compared to non-VHBA counterparts. The higher incorrect rate in the TAP incompatible task indicates that VHBA subjects have poorer ability to manage interference in stimulus-reaction incompatibility. These subjects experience difficulty in recognizing the pointing direction of an arrow when its direction differs from its location of appearance on the screen. Counterintuitively, the TAP compatible incorrect rate of VHBA subjects was significantly lower, suggesting that VHBA individuals could better recognize an arrow that was pointing in the same direction as its appearance. On the other hand, VLBA old adults demonstrated higher correct rates in recognizing target real words from distractor pseudo words in the WST task, compared to non-VLBA old individuals. Previous studies have confirmed that chronological age affects neurocognitive functions such as working memory and attention [32, 2]. We observed that WST correct rates, as well as TAP compatible and incompatible incorrect rates, showed no significant differences between subjects with young and old chronological age. These findings suggest that high or low brain age, irrespective of chronological age, could be linked to neurocognitive functions.

We found that most young adults with psychological conditions and addictions manifested high brain age. MRI studies have found that excessive alcohol consumption is associated with damage to the frontal lobe [33, 34], including functional connectivity and EEG activity changes [35]. Long-term cannabis consumption was also linked to reduced gray matter volume in the temporal and frontal cortices in MRI study [36]. Similar to the aging brain, excessive alcohol and cannabis consumption affect working memory, attention, and verbal functions [37, 38, 39]. Phobia patients demonstrate abnormalities in the frontal and visual cortices [40, 41], while depressed patients have impaired frontal and temporal cortices [42, 43]. The grey matter volume of the medial prefrontal cortex (MPFC), which is responsible for emotion and cognitive functions, was also found to be reduced in depressed individuals’ brains [44]. Similarly, depressed individuals exhibit poorer memory, attention, and executive functions compared to healthy subjects [45, 46, 47]. The frontal cortex, which is the main brain area affected by these psychological conditions and addictions, is also the area most affected by aging [3, 48]. This explains the relationship between the above-mentioned mental conditions and brain age. In contrast to MRI-based brain age studies that typically highlight structural changes as key markers of aging, our study based on functional connectivity measured by EEG could provide a complementary functional perspective to structural MRI, combining the findings from both modalities offer a more comprehensive depiction of brain aging.

The results of this study indicate that VHBA adults have lower frontocentral connectivity and nodal degrees in wide, theta, and low gamma frequency ranges. This is in line with previous studies that reported decreases in frontocentral theta power are related to normal aging and cognitive functions such as memory and attention [3, 48, 2, 49]. The involvement of central motor areas and low gamma band is likely due to the normal degradation of motor functions in the elderly [50, 51]. On the other hand, VLBA individuals have lower prefrontal connections and higher frontal degrees, especially in wide, delta, theta, and alpha frequency ranges. We hypothesize that VLBA individuals can maintain localized brain functions (higher degrees) with lower demands on prefrontal connections. In addition to theta activity, memory-consolidating delta and attention-modulating alpha activity were also found to change with age [52, 53, 54]. Subsequent analysis found that delta frontocentral and low gamma prefrontal regions were significantly correlated with brain age but not chronological age, further supporting the relationship of these regions with brain age. These results are consistent with the findings of our ablation study, which showed that the combination of frontal and central regions provided the best prediction of brain age.

In conclusion, this study linked brain age predicted from EEG functional connectivity with cognitive and clinical manifestations. Brain age was found to be related to memory, attention, substance addictions, phobia, and depression. These mental conditions, together with normal aging, mainly affect the frontal brain regions. This leads to improved predictability of brain age using frontal activity. There are several limitations in this study. This study could not completely eliminate the fundamental limitation of regressing toward the mean [55, 56], where the ages of young adults were overestimated and the ages of old individuals were underestimated. However, our results, which found significant differences between VHBA/VLBA and baseline subjects, could suggest the robustness of our findings. The relatively high MAE in this study was likely due to the stratification of ages in 5-year ranges, which could be addressed in future studies by obtaining the exact age of each subject. Besides, this study investigated the correlation between functional connectivity and brain age, subsequently, longitudinal study could be conducted to study the respective causal effect. The performance of regression models, such as support vector machines, convolutional neural networks, transformer networks, and recurrent neural networks, could also be evaluated with larger dataset in future studies.

